# Shifting gradients of macroscale cortical organization mark the transition from childhood to adolescence

**DOI:** 10.1101/2020.11.17.385260

**Authors:** Hao-Ming Dong, Daniel S. Margulies, Xi-Nian Zuo, Avram J. Holmes

## Abstract

The transition from childhood to adolescence is marked by pronounced shifts in brain structure and function that coincide with the development of physical, cognitive, and social abilities. Prior work in adult populations has characterized the topographical organization of cortex, revealing macroscale functional gradients that extend from unimodal (somato/motor and visual) regions through the cortical association areas that underpin complex cognition in humans. However, the presence of these core functional gradients across development as well as their maturational course have yet to be established. Here, leveraging 378 resting-state fMRI scans from 190 healthy individuals aged 6-17 years, we demonstrate that the transition from childhood to adolescence is reflected in the gradual maturation of gradient patterns across the cortical sheet. In children, the overarching organizational gradient is anchored within unimodal cortex, between somato/motor and visual territories. Conversely, in adolescence the principal gradient of connectivity transitions into an adult-like spatial framework, with the default network at the opposite end of a spectrum from primary sensory and motor regions. The observed gradient transitions are gradually refined with age, reaching a sharp inflection point in 13- and 14-year-olds. Functional maturation was nonuniformly distributed across cortical networks. Unimodal networks reached their mature positions early in development, while association regions, in particular medial prefrontal cortex, reached a later peak during adolescence. These data reveal age-dependent changes in the macroscale organization of cortex and suggest the scheduled maturation of functional gradient patterns may be critically important for understanding how cognitive and behavioral capabilities are refined across development.

**Significance:** Human abilities and behavior change dramatically across development, emerging from a cascade of hierarchical changes in brain circuitry. Here, we describe age-dependent shifts in the macroscale functional organization of cortex in childhood and adolescence. The characterization of functional connectivity patterns in children revealed an overarching organizational framework anchored within unimodal cortex, between somato/motor and visual regions. Conversely, in adolescents we observed a transition into an adult-like gradient that situates the default network at the opposite end of a spectrum from primary sensory and motor regions. This spatial framework emerged gradually with age, reaching a sharp inflection point at the transition from childhood to adolescence. These data reveal the presence of a developmental change from a functional motif first dominated by the distinction between sensory and motor systems, and then balanced through interactions with later-maturing aspects of association cortex that support more abstract cognitive functions.

## Introduction

The cerebral cortex is comprised of a complex web of large-scale networks that are central to its information-processing capabilities (1, 2). Although the topographic organization of this distributed network architecture has been characterized in adulthood (3–6), there is a growing consensus that macroscale patterns of functional connectivity are not static across the lifespan (7–12). In particular, the transformative brain changes occurring throughout childhood and adolescence are critical for supporting the emergence and development of physical, cognitive and social abilities (13–17). Yet despite clear evidence that patterns of brain organization from neural circuits through large-scale cortical networks relate to behavior (18–20), the age-dependent changes that underpin hierarchical shifts across the functional connectome have yet to be systematically investigated.

Technical advances in the field of connectomics have provided researchers with tools to map the topographic organization of large-scale distributed networks in the human brain (3, 5, 6). The cortical sheet is comprised of areal units, traditionally defined through their embryological origins, cytoarchitecture, and evoked functions (21–23). Critically, the brain is a multi-scale system and these areal parcels are embedded within segregated processing streams and corresponding networks that are evident through both anatomical projections and patterns of coherent neural activity at rest (2, 24, 25). This converging evidence suggests the presence of a broad division separating the unimodal somatosensory/motor (somato/motor) and visual territories that form domain-specific hierarchical connections (22) and the heteromodal association areas that integrate long-distance projections from widely distributed sets of brain systems (26). However, the distinction between unimodal and heteromodal cortex is not reflected in abrupt transitions along the cortical surface. Recent work has revealed that this macroscale property of brain organization is evident in the presence of a principal functional gradient that situates discrete large-scale networks along a continuous spectrum, extending from the unimodal systems that underpin perception and action through the association territories implicated in more abstract cognitive functions (12, 27, 28).

Cortical gradients capture the topography of large-scale networks, offering a complementary approach to standard brain parcellation techniques. Here, rather than marking discrete functional boundaries, spatial variation across the cortex is examined continuously along overlapping organizing axes (27, 29). In both humans and non-human primates, the association cortex end of the principal gradient is anchored within the default network, including portions of ventral and dorsal medial prefrontal, posterior/retrosplenial, and inferior parietal cortex (30–32). The default network acts as a hub that integrates representational information across the cortex (1, 6). As such, it constitutes a functional system hypothesized to underpin self-referential processing and core aspects of mental simulation (33). It has recently been proposed that the default network sits at the apex of cortical hierarchy, situated as the most distant association network from sensory cortex (34). This feature of brain organization that may reduce the strong constraints of sensory/motor input on default network function. Suggesting cognitive abilities are closely associated with these macroscale gradients, the spatial framework evident in profiles of intrinsic brain activity mirrors the patterns of functional specialization and flexibility emerging through extrinsic (task-evoked) studies of the human brain (19). These functional gradients have been well characterized in adult populations, providing a framework for describing the integration of local processing streams throughout cortical (27) and subcortical systems (29). However, the presence of this overarching organizational structure across development as well as its maturational course have yet to be established.

Human brain development is influenced by a complex series of dynamic processes across biological systems, ranging from shifting profiles of gene expression (35) to hierarchical changes in brain structure and function (14). Although still in its early stages, work in this area has characterized several core organizational principles underlying large-scale network development (36–40). In human and non-human primates, for instance, there is an earlier maturational plateau in gray matter (41–43) as well as in synaptic formation and subsequent pruning within unimodal sensory/motor and subcortical territories relative to aspects of association cortex (42, 44). This developmental sequence is reflected in patterns of network activity. For example, brain functions in infancy are characterized by the predominance of short-range connectivity (45, 46) which gradually transitions through childhood and adolescence as long-range network connections becomes increasingly evident (11, 47–52). Collectively, these results suggest that brain development may entail a shift from a sensory segregation (unimodal gradient) to a distributed global integration (association gradient) across childhood and adolescence. Whole-brain connectome-wide neurodevelopmental studies have identified patterns of functional network organization that are predictive of behavioral traits (53) and associated with mental health outcomes in adolescence and young adulthood (7). However, to date there have been few opportunities to directly explore the age-dependent maturation of functional gradients across cortex. The characterization of age-dependent changes in the macroscale organization of the human brain would provide a tremendous opportunity to understand how connectome development shapes the evolving expression of individual differences in behavior across health and disease.

In examining age-dependent shifts in the macroscale functional organization of cortex we applied the dimensionality reduction approach of diffusion map embedding (27, 54) to resting-state fMRI data to extract a global framework that accounts for the dominant connectome-level connectivity patterns in a population of children and adolescents. The resulting components, or gradients, reflect the distillation of complex patterns of local and global functional connectivity across the connectome into simple and interpretable spatial architectures. This approach allowed us to establish the extent to which functional maturation is nonuniformly distributed across cortical networks. In doing so our analyses revealed the presence of a developmental change from a functional motif first dominated by the distinction between sensory and motor systems, and then balanced through interactions with later-maturing connectivity within association cortex.

## Results

### The dominant connectivity gradient transitions from a unimodal network architecture in children to association cortex in adolescents

We first decomposed the functional connectivity matrix into components that capture the maximum variance separately for children and adolescents. Functional connectivity matrices consisted of 20,484 vertices across the cortical sheet derived in data from the Chinese Color Nest Project (CCNP) (56–58). Initial age-related differences in gradient architecture were evaluated by dividing participants’ scans into child (n=202) and adolescent groups (n=176). Consistent with prior work (27), diffusion map embedding (55) was used to decompose participant connectivity matrices, reducing data dimensionality through the nonlinear projection of the vertices into an embedding space. The resulting functional components reflect divergent spatial patterns of connectivity across cortex, ordered by the variance explained in the initial functional connectivity matrix (Supplemental Table 1). Within each component, similar connectivity patterns result in similar values or ‘connectopies’ (28), here referred to as ‘gradients’ (27). Applied in this manner, diffusion map embedding can distill complex patterns of local and global functional connectivity into simple functional components with interpretable spatial architectures (27, 28, 54). The resulting gradients, are unitless and reflect the position of vertices along an associated embedding axis that captures the primary differences in vertex-level connectivity patterns.

The first two gradients, accounting for the greatest variance in functional connectivity, differed in their spatial organization between children and adolescents (Figure 1A; Supplemental Figure 1). The peak gradient values in children were situated at the central and calcarine sulci, revealing a spectrum differentiating somato/motor and auditory cortex from the visual system. The first gradient in children closely resembles the second gradient previously identified in adults (27) as well as the second gradient identified in adolescents in the present analyses (Figure 2). Conversely, in the adolescent group, the first gradient resembled the principal gradient in adults (27). Although one end of the principal gradient of connectivity in adolescents was anchored in somato/motor and auditory cortex, the regions at the other end encompassed broad swaths of association cortex, including portions of ventral and dorsal medial prefrontal, posteromedial/retrosplenial, and inferior parietal cortex — a spatial pattern that corresponds with the canonical default network (33, 34). Scatter plots further characterize the switch between the two dominant gradient patterns that accompany the transition from childhood through adolescence and into adulthood (Figure 1B).

**Figure 1:**
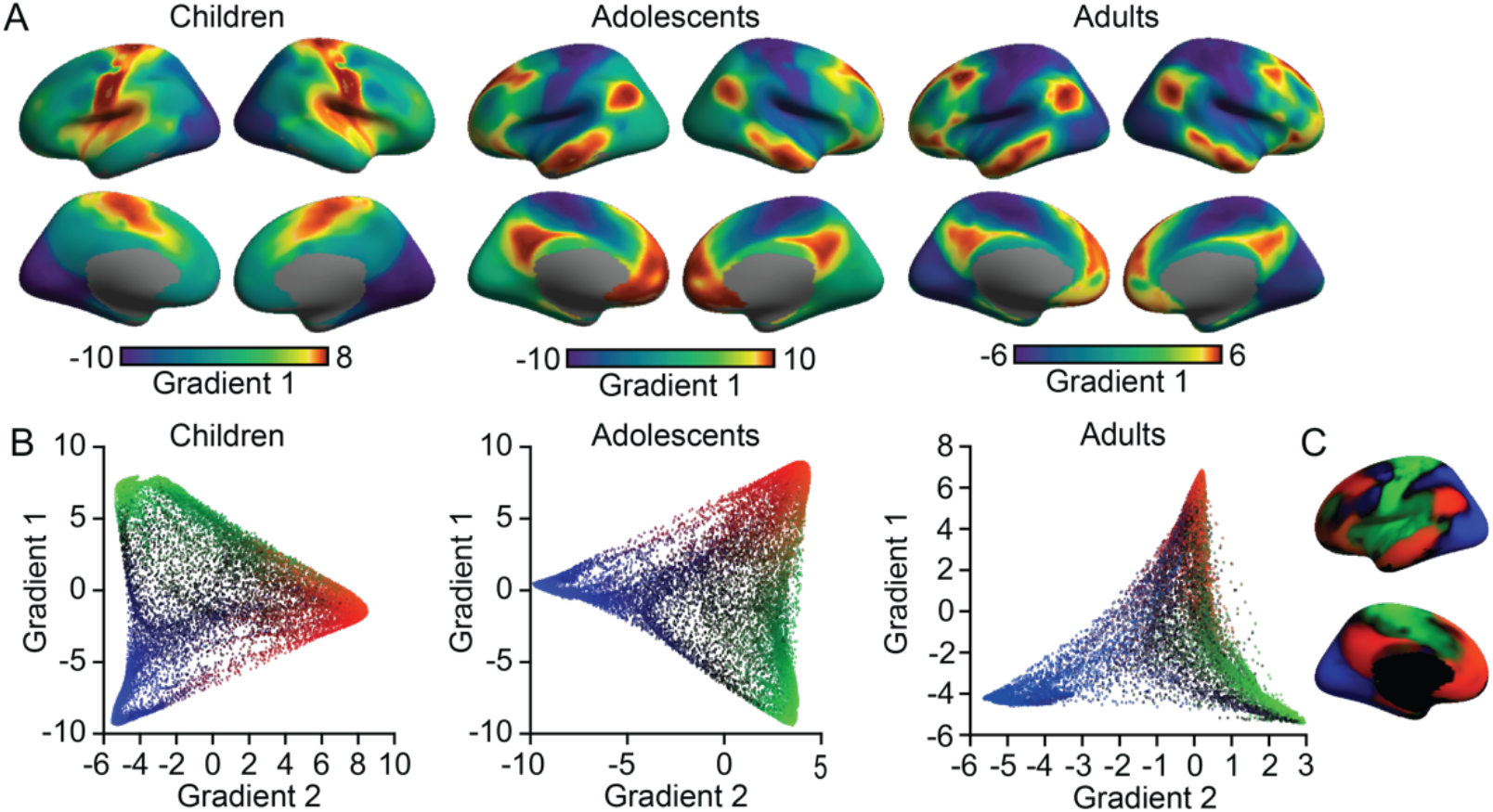
Functional connectivity gradients differ between childhood and adolescence. (A) The principal gradient of connectivity in children (left panel) peaks within unimodal networks, separating somato/motor and auditory cortex (red) from visual cortex (blue). Conversely, in adolescents (middle panel) the principal gradient reveals adult-like organization (right panel, data from Margulies et al., (27)), transitioning from unimodal to association cortex (red). The proximity of colors within each age group indicate the similarity of connectivity patterns across the cortex. Scale bar reflects Z transformed principal gradient values derived from connectivity matrices using diffusion map embedding (55). (B) Scatter plots of the first two connectivity gradients in children, adolescents, and adults. A clear switch within the two gradients accounting for the maximum variance in connectivity is evident between the children and adolescents. (C) Colors from scatter plot (B) are presented on of the cortical surface. In children, gradient 1 separates somato/motor (green) and visual regions (blue), while gradient 2 distinguishes unimodal from association cortex (red). These two gradient patterns are flipped in adolescents.

**Figure 2:**
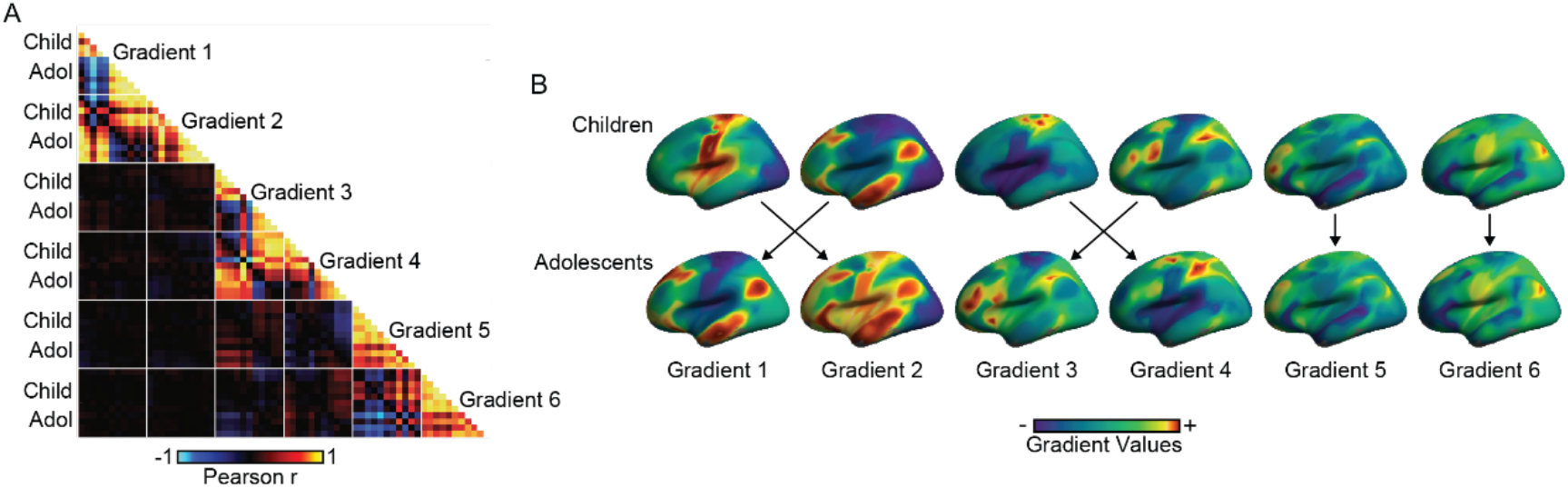
Evidence for both common and distinct gradient architectures across childhood and adolescence. (A) Similarity matrix presents the vertex-level Pearson correlations for the first 6 gradients across participants grouped at 1-year age intervals. Within-age group correlations for each gradient fall along the center diagonal. Between-age group and cross-gradient correlations are plotted away from the diagonal and reveal both positive (red) and negative (blue) relationships. Participants under 12-years-old are labeled as children. Here, associated rows and columns correspond to 6-7, 8, 9, 10, and 11-year-old groups. Adolescent associated rows and columns correspond to groups of 12, 13, 14, 15, 16, and 17-year-old participants. White lines represent gradient boundaries. (B) Gradient profile maps rendered on cortical surface reveal transitions between the first 4 gradients across childhood and adolescence. In contrast, the architectures in gradient 5 and 6 remained consistent across groups. Black arrows denote maximal vertex-level correlations across age groups for each gradient. Child, children; Adol, adolescents.

### The transition from childhood to adolescence is marked by both dynamic and stable functional gradient architectures

Previous studies of the macroscale functional organization of the cerebral cortex in adulthood have largely focused on the principal gradient in connectivity in isolation. Having established the presence of distinct gradient topographies in childhood and adolescence, we next examined the consistency of the first six gradients across development (Figure 2) accounting for the bulk of the functional connectivity variance in children (percent variance: 73.2) and adolescents (percent variance: 72.2). Correlations across 1-year interval age groups were also calculated (Figure 2A; see Supplementary Data for full set of gradient comparisons).

Human brain development is characterized by the early maturation of unimodal visual and somato/motor cortex, the subsequent refinement of multimodal association areas (41, 59), and the gradual integration of the visual system into a global processing hierarchy (12). Consistent with these developmental trajectories the primary gradient of connectivity in children most closely resembled the secondary gradient in adolescents (Pearson’s r=0.80). Further, the secondary gradient of connectivity in children most closely resembled the primary gradient in adolescents (r=0.80). The observed relationships within the first two connectivity gradients across development are consistent with a gradual transition in which sensory/motor regions accounting for the most variance in connectivity in childhood are replaced by a principal gradient in connectivity that spans primary/unimodal and association cortex in adolescence. Of note, the secondary gradient in adolescents exhibited a hybrid structure containing both within-unimodal differentiation and contrast between unimodal and association regions. In adults, the component accounting for the second-most variance in connectivity differentiates regions solely within the unimodal end of the principal gradient (27). While the present analyses are consistent with the initial emergence of an adult-like second gradient in adolescence, future work should further characterize the trajectories of gradient organization across the lifespan (12).

The developmental transition from a unimodal segregation to transmodal integration organizing framework was also evident in the third and fourth gradients. Here, the third gradient in children was anchored in somato/motor cortex, closely resembling the fourth gradient in adolescents (r=0.82). Conversely, the fourth gradient in children, reflecting a network architecture contrasting frontoparietal and somato/motor systems, mirrored the third gradient in adolescents (r=0.85). This pattern of adolescent brain function that follows initial reports of gradient topographies in adults (27). As a whole, these cross-gradient relationships are consistent with a broad developmental transition throughout cortex in the functional variance accounted for by somato/motor and visual regions to a more distributed connectivity structure that encompasses higher-order association areas. Importantly, the observed developmental transitions were not ubiquitous across all gradients. Highlighting the presence of stable features of the macroscale organization of cortex, the fifth and sixth gradients were spatially consistent between the child and adolescent groups (rs≥0.90). Moving forward, the analysis of stable and dynamic connectivity structures across development may help determine the unique neurocognitive trajectories that emerge across childhood and adolescence.

### Principal gradients of connectivity gradually transition from a unimodal to association cortex architecture across development

The use of categories, such as child or adolescent, can reveal stable gradient architectures shared within a broad developmental stage. However, many age-specific properties of brain function are lost when central tendencies are examined across large groups, potentially masking subtle shifts associated with the gradual process of neurodevelopment. As one example, the current observation of a hybrid unimodal and association cortex structure for the secondary gradient in adolescents could emerge through at least two potential routes. First, given the age ranges included within the child (6-12 years of age) and adolescent (12-18 years of age) groups, within-population heterogeneity may obscure or blur the scheduled maturation of functional gradient patterns. An alternate, but not mutually exclusive, possibility is that adult-like functional integration may emerge later in adolescence, delaying the formation of fully distinct gradient patterns across development. To examine these hypotheses, we rederived the gradient architectures in participants at 1-year age intervals.

The principal and secondary gradients of connectivity for each age group are displayed along the lateral surface of the left and right hemispheres in Figure 3. For the first gradient, participants below 12 years of age displayed unimodal dominant profiles. An abrupt transition was evident in the 12-year-old participants, from this point forward all individuals exhibited an adult-like gradient organization with default network anchored at the opposite end of a spectrum from primary sensory and motor regions. The second gradient followed an inverse but smoother transition profile across development. Here, the extreme ends of gradient 2 reflect a mixture of unimodal and association cortex until approximately 14 years of age. At this point, gradient values within association cortex gradually decrease, while the contrast between somato/motor and visual regions becomes more apparent. We quantified this trend by extracting the gradient values (standard z-scores) within the default network across each 1-year age interval (Figure 3B). Across both gradients, age 12 reflected a transition point where the associated default network gradient values began to reverse. For the first gradient, the relative gradient values within the default network fluctuated around 0 prior to 12 years of age. These data suggest a relatively minimal influence of the default network on this gradient pattern until adolescence. The reversed pattern in the second gradient is consistent with a gradual developmental convergence towards a global network structure where association cortex becomes further differentiated and integration across distinct systems becomes the dominant feature of cortical organization. Further analysis establishes that the gradient value within default network reaches peak later in adolescence (Figure 3C). Radar plots reveal the point at which network gradient values peak during development across each of the first six gradients (Supplemental Figure 2).

**Figure 3:**
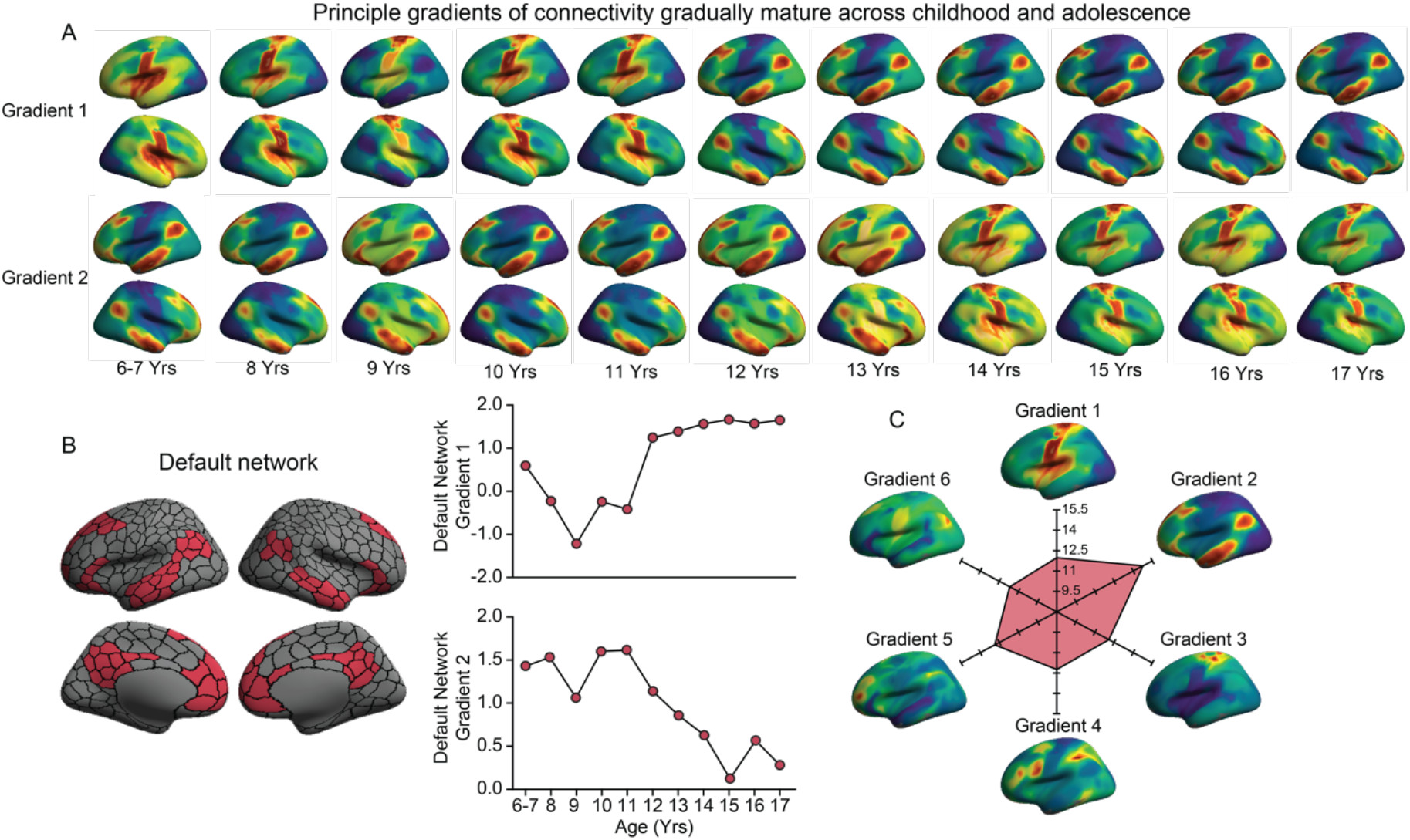
Functional connectivity gradients gradually mature across childhood and adolescence. (A) Participants are divided into groups at 1-year age intervals. Gradient values are displayed on the lateral surface of the left and right hemispheres where the proximity of colors within each age group indicates the similarity of connectivity patterns across the cortex. The principal gradient (gradient 1), which accounts for the greatest variance in connectivity, segregates unimodal regions in childhood. Across development a gradual transition was observed in gradient 1 reflecting the shift to an adult-like architecture after 12 years of age, at one end anchored by sensory and motor regions and at the other end by association cortex. Gradient 2 surface maps reveal the inverse transitional profile across development. Here, the extreme ends of gradient 2 reflect a mixture of unimodal and association cortex until 14 years old, at which point gradient values within association cortex decrease, while the segregation between somato/motor and visual regions becomes more prominent. (B) Red parcels denote regions within the default network. Black lines denote parcel and network boundaries based on the Yeo et al., (3) 7-network solution averaged across the 400-parcel functional atlas of Schaefer and colleagues (60). Graphs display average gradient 1 and gradient 2 values within the default network for each 1-year bin. (C) Radar plot displays the age at which the default network gradient values peak during development across each of the first six gradients. Radar plots for the remaining large-scale networks are available in Supplementary Figure 2.

We next examined the extent to which the first two gradients in childhood and adolescence capture the macroscale layout of canonical large-scale functional networks in adulthood. Here, we examined the Yeo et al. (3) 7-network solution averaged across vertices. Gradient values of all seven functional networks are summarized using boxplots in Figure 4. In line with prior work in adult populations (27), networks were not randomly distributed along gradients in either children or adolescents. Rather, cortical parcels from the same network tended to cluster at similar positions (Figure 4B). In children, the somato/motor and visual systems were situated at the extremes along the first gradient. The second gradient in children resembled a transmodal integration dominant gradient profile, previously reported in adults to follow the spatial constraints of cortical anatomy (27). Here, the default network occupied one extreme position along the second gradient and was maximally separated from the visual system from the perspective of functional connectivity. Conversely, in adolescents, both the first and second gradients consisted of a mixture of unimodal and association cortex architectures. Here, the default network occupied an extreme end of both gradients. Despite this commonality, the first two adolescent gradients were still distinguishable through the distinct integration of somato/motor and visual regions. The first gradient of adolescents was characterized by a heteromodal integration structure, where somato/motor regions were anchored at one end and the visual system was situated at the center of the spectrum. This contrasts with the second gradient in adolescents where, although the default network occupied one extreme, somato/motor regions also approached the same gradient terminal with visual regions occupying another end (Figure 4B). This profile suggests a secondary gradient structure approaching a unimodal segregation organizational scheme, but that is still in progress during adolescence. Collectively, these results demonstrate clear transitions between the principal and secondary gradients of connectivity across childhood and adolescence, providing a framework for the future study of the spatial ordering of large-scale networks during this critical stage for the development of association cortex.

**Figure 4:**
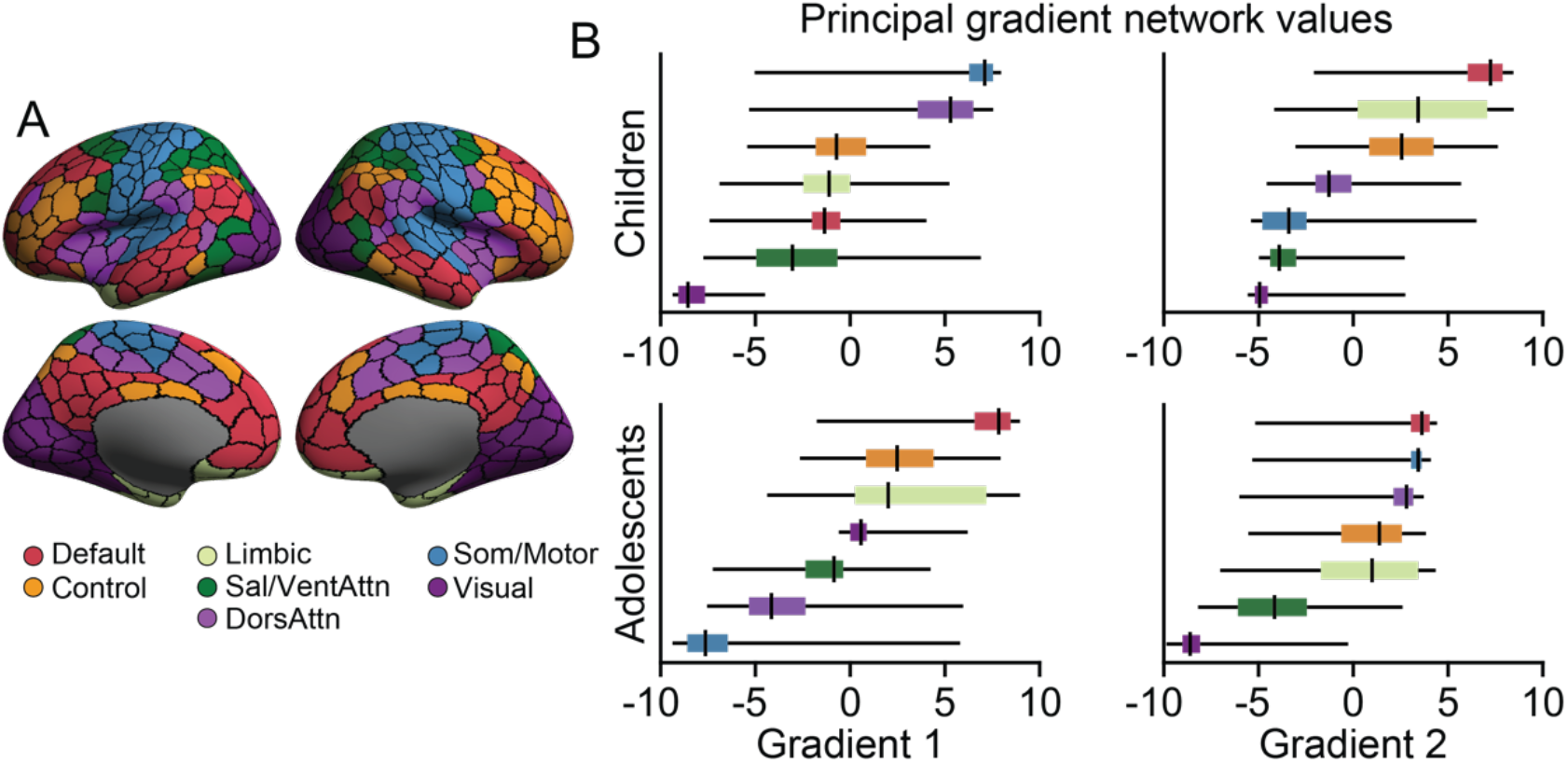
The default network occupies the extreme end of both principal gradients in adolescence, but not childhood. (A) Vertex level data were summarized based on the Yeo et al., (3) 7-networks. (B) The boxplots reflect the gradient 1 and gradient 2 values averaged within 7 networks in children and adolescents, ordered by median value. In children, unimodal regions are situated at the extremes along the gradient 1 while the default network assumed this position in gradient 2. Conversely, in adolescents the default network occupied the extreme position along the first two dominant gradient patterns.

### Functional maturation is nonuniform within heteromodal and unimodal cortex

Our findings detailed above reveal age-dependent changes in the macroscale topographic organization of cortex, highlighting the transition from a functional motif first dominated by sensory and motor circuits in childhood into one balanced through interactions with later-maturing association cortex in adolescence. Yet, an important unanswered question remains the extent to which the observed developmental cascades are spatially uniform within large expanses of the cerebral cortex or are nonuniformly distributed at a more granular level both across and within discrete cortical networks.

Here, gradient values were normalized across all surface vertices. Within each gradient map, the age at which the standard gradient value reaches its peak was assigned to each vertex as its ‘maturation age.’ The average of the six maturation maps was obtained to represent the process of functional maturation (Figure 5A). Critically, functional maturation was not randomly distributed across cortex, instead the spatial topography of the maturation map broadly adhered to parcellation borders separating unimodal and association cortex (3, 60). Visual cortex and somato/motor regions mature earlier in childhood along with portions of anterior insula. The subsequent age at which gradient values peak during development was not uniform within association cortex. Rather, the early unimodal segregation-dominant profile is followed by subsequent gradient peaks within frontoparietal control and attention networks, with limbic and default networks maturing later in adolescence (Figure 5B). In particular, core default network regions including medial prefrontal cortex, parahippocampal gyrus, superior temporal sulcus and inferior frontal gyrus, were observed to mature latest in adolescence. The maturation map also shows sensitivity in distinguishing functionally heterogenous, yet spatially contiguous cortical territories. As one example, the inferior and superior parietal lobules are spatial adjacent, but the former is functionally coupled to the default network while the latter comprises part of the frontoparietal network. Consistent with the complex functional organization of the parietal lobe, gradient maps revealed tightly interdigitated but differentiated patterns of functional development.

**Figure 5:**
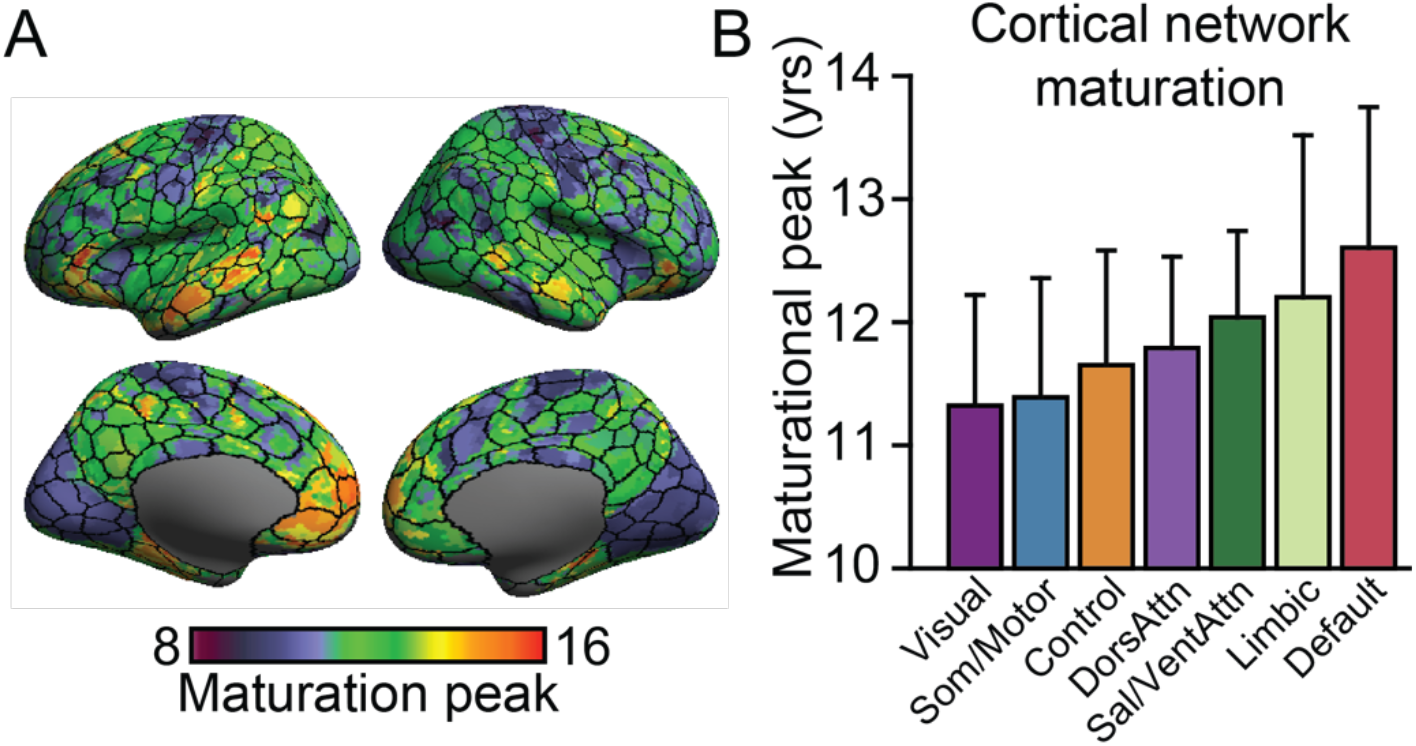
Functional maturation is nonuniformly distributed across cortical networks. (A) Surface maps display vertex-level functional maturation with the 400 parcel functional atlas (borders from Schaefer and colleagues (60) are overlaid). Scale bar reflects the ages at which gradient values peak during development. The topography of functional maturation broadly adheres to parcellation borders separating unimodal and association cortex. (B) Age at which the six gradient values peak during development was averaged and plotted within the boundaries of each network. Error bars denote standard deviation. Maturational age broadly follows the cognitive processing hierarchy. Somato/motor and visual networks mature in childhood while association regions, in particular medial prefrontal aspects of default and limbic networks, peak later during adolescence.

## Discussion

The cerebral cortex is tiled with networks of distinct cortical areas, often delineated by embryology, cytoarchitecture, and evoked functions, broadly linked through continuous gradients of connectivity. In adults, this complex organizational profile can be represented in a low-dimensional space anchored, at one end, by unimodal regions supporting primary sensory/motor functions and at the other end, by association cortex (27). However, human abilities and behavior change dramatically across development and the maturational course of these core functional gradients have yet to be established. Here, in a population of school-age children and adolescents, we utilized diffusion map embedding (27, 54) to reveal age-dependent shifts in the macroscale organization of cortex. These analyses demonstrate the presence of a gradual developmental change from a functional connectivity motif in childhood, first dominated by the distinction between unimodal systems, and later balanced in adolescence through the maturation of a functional hierarchy of integration that supports more abstract cognitive functions.

Human brain development is marked by a complex series of dynamic processes including hierarchical changes in brain systems that subserve executive functions, affect regulation, and social abilities (14, 16, 17). Of particular relevance for our current analyses, work in animal models has revealed that while synaptogenesis and pruning in unimodal cortex occurs earlier in life (61), association cortex is characterized by a protracted period of synaptic development that continues into adolescence (62, 63). This staged developmental pattern is consistent with post-mortem work in humans suggesting that by age 12 synaptic pruning within auditory cortex is complete, a process that extends through mid-adolescence within the middle frontal gyrus (42). Along the same lines, age-related change is evident in the staggered maturation of cortical gray matter across development where association cortex matures after unimodal regions (41), at least in part reflecting corresponding shifts in the degree of intracortical myelination (64). The dynamic progression of development from primary sensory and motor systems through networks supporting flexible and abstract cognition likely has a profound impact on the computational landscape of the brain, although studies of brain function have often focused on regional approaches or discrete circuits (but see (48, 49, 65, 66)). Here, we extend upon this literature to characterize the macroscale functional organization of cortex in children and adolescents. Consistent with the development of regional milestones, our results highlight the gradual maturation of broad connectome-level gradients across development, reaching a tipping point around age 13. In children, the bulk of the functional variance was accounted for by a unimodal architecture anchored between somato/motor and visual regions. Conversely, in adolescence the dominant functional gradient gradually transitioned into an adult-like spatial framework, with the default network at the opposite end of a spectrum from primary sensory and motor regions. Of note, the current sample of adolescents exhibited less differentiation between transmodal and visual regions, relative to somato/motor. These data suggest the likely continued refinement of the visual system within the global connectivity structure through young adulthood (12).

Although the pattern of cortical maturation was broadly consistent at the network-level within unimodal and heteromodal systems, we observed substantial spatial variability in the sequence of gradient development across the cortex. Cortical gray matter develops in a staggered regional manner, with the territories supporting primary functions developing in advance of regions involved in complex and integrative tasks (41). Here, functional maturation broadly followed the cognitive processing hierarchy. Consistent with prior work, somato/motor and visual networks reached a maturational peak in childhood while association regions, in particular temporal and medial prefrontal components of default and limbic networks, peaked later during adolescence. The maturation of default network typically progresses along a local to distributed pattern. Although the topological profile of the default network has been found as early as the first year after birth (46), local connections are dominant between network hubs during childhood, while distant connectivity is fragmented (49). It is not until late adolescent and early adulthood that dense long-range connections emerge (9, 11). In the present analyses, an adult-like default network-anchored gradient was evident during early childhood accounting for ~10-14 precent of the observed functional variance, relative to ~34-39 precent for the unimodal-centered gradient. Across development we observed a gradual transition in the dominant topographic organization of large-scale connectivity, as the amount of variance accounted for by the unimodal spectrum decreased relative to the transmodal integration dominant gradient architecture. These data are consistent with prior regional and circuit focused analyses demonstrating the staggered development of the amygdala and the medial prefrontal cortex across childhood and adolescence (14) and associated shifts in responses to emotionally evocative stimuli (67–69). Here, the use of diffusion map embedding allowed for the examination of the hierarchical development of multiple networks across childhood and adolescence. Suggesting a broad process of association cortex maturation beyond the conical default network, age-dependent changes in brain function were not limited to the primary and secondary gradients. For example, across the third and fourth gradients the increased functional importance of a differentiation of the frontoparietal network became evident during the transition into adolescence.

The present results suggest the age-dependent refinement of a principal gradient that situates discrete large-scale cortical networks along a continuous spectrum, beginning within unimodal input–output systems and ending within the default network. Of note, the reported functional connectivity patterns reflect group averages across subjects. Accordingly, the amount of variance explained within specific networks may be due, at least in part, to associated patterns of inter-subject variability. Critically however, the age-dependent transition in gradient architectures allows us to infer the complex reorganization of hierarchical network relationships across development. It has been proposed that the spatial arrangement of areal parcels across the cortical sheet from motor and sensory regions, through attention and cognitive control related territories, finally terminating at the default network provides anatomical constraints on information processing (27, 34). Given the core role of association cortex — in particular the default network — in a host of traits and abilities (70), including affect regulation and social functioning (71), as well as increased vulnerability for association cortex mediated pathologies in adolescence (e.g., affective illnesses; (72)), future work should examine possible relationships linking large-scale gradient transitions with symptom burden across development.

The present data are consistent with prior work suggesting a repeating evolutionary and developmental motif embedded within the macroscale properties of human brain organization. Although the fundamental organization of the sensory and motor areas that comprise unimodal cortex emerged early in vertebrate evolution (73), as brain sizes have increased in primates, association cortex has disproportionately expanded (74, 75). This evolutionary sequence of differential cortical scaling is mirrored across developmental periods, as reflected in the staggered embryonic development of the cells that colonize the cortical sheet (76), the formation of subcortical-cortical connections in infancy (8), and the present analyses revealing the gradual refinement of large-scale gradients across childhood and adolescence (9, 49). For instance, this is evident in the regulatory role of cortico-thalamic connections in the formation and maintenance of functional assemblies across cortex (77). Reflecting the presence of a scheduled maturation in the development of thalamic projections to unimodal and association networks, functional connectivity between the thalamus and sensory/motor cortex is evident in human neonates, while connectivity between the thalamus and association cortex does not emerge until the first year of life (78). Although speculative, such phenomenon hint at the possible presence of a universal rule of maturational order where the same organizing heuristics may be invoked multiple times across the lifespan, guiding the development of cerebral cortex. An important question then arises as to what factors may serve as the early foundation for the proto-organization of cortex across development and influence subsequent periods of functional refinement during the transition from childhood to adolescence at the onset of puberty.

How the functional architecture of the cerebral cortex forms and is refined throughout development is a central and challenging question across the neurosciences. The present results demonstrate age-dependent changes in the topographic organization of cortex, transitioning from a gradient differentiating sensory/motor modalities into a functional motif which is later balanced through an increased emphasis on transmodal integration within association cortex. While a broad process of association cortex maturation was evidence across default and frontoparietal networks, we observed substantial spatial variability in developmental timing indicting that the maturational age of cortical regions was not exclusively driven by spatial proximity. Although our understanding of the functional consequences of gradient refinement across development is incomplete with respect to their roles in behavior across health and disease, the stereotyped progression from unimodal through association cortex architectures nominates candidates for subsequent functional experiments. Whether this phenomenon reflects a universal rule of maturational order in humans, remains a hypothesis to be tested in future work.

## METHODS

### Chinese Color Nest Project

The Chinese Color Nest Project (CCNP) is a five-year accelerated longitudinal study across the human life span (56–58). At baseline of the recruitment, CCNP ‘growing up in China’ invited a total of 198 school age participants from the Chinese Han population in Chongqing. This accelerated design is particularly valuable for the group-average developmental analysis employed in the present work because each individual was invited to undergo three magnetic resonance imaging (MRI) scans at an interval of 15 months (balancing season effects) and the longitudinal scans of an individual participant can be included in different age groups. We thus generated a dataset of 176 adolescents and 202 typically developing children. Any participant with a history of neurological or mental disorder, family history of such disorders, organic brain diseases, physical contraindication to MRI scanning, a total Child Behavior Checklist (CBCL) T-score higher than 70, or a Wechsler Intelligence Scale for Children IQ standard score lower than 80 were excluded from further analysis (56). All MRI data was obtained with a Siemens Trio 3.0T scanner at the Faculty of Psychology, Southwest University in Chongqing. For each visit, the scanning order was as follows: resting-state fMRI scan (7min 45s), T1 MP-RAGE scan (8min 19s), resting-state fMRI scan (7min 45s), giving a total of 15min 30s resting-state fMRI scanning. Details regarding participant recruitment and characterization, as well as the demographic characteristics of the sample are available in Dong et al. (56). The resting-state scans were acquired with an echo-planar imaging (EPI) sequence using following parameters: flip angle=80°, FOV=216mm, matrix=72×72, slice thickness/gap=3.0/0.33 mm, TR/TE=2500/30ms, slice orientation: sagittal, acquisition direction: interleaved ascending, number of measurements=184, scanning time lasted for 7min 45s. The reported experiments were approved by the Institutional Review Board from Institute of Psychology, Chinese Academy of Sciences. All participants and their parents/guardians gave written informed consent before participating in the study.

### MRI data preprocessing

Anatomical T1 images were visually inspected to exclude individuals with substantial head motion and structural abnormalities. Next, T1 images were fed into the volBrain pipeline (http://volbrain.upv.es) (79) for noise removal, bias correction, intensity normalization and brain extraction. All brain extractions underwent visual inspection to ensure tissue integrity. After initial quality checks, T1 images were passed into the Connectome Computation System (CCS) (80) for surface-based analyses. CCS pipeline is designed for preprocessing multimodal MRI datasets and integrating several publicly available software such as SPM (81), FSL (82), AFNI (83) and FreeSurfer (84). For resting-state fMRI data, preprocessing included a series of steps common to intrinsic functional connectivity analyses: (1) dropping the first 10s (4 TRs) for the equilibrium of the magnetic field; (2) head motion correction; (3) slicing timing; (4) de-spiking for the time series; (5) estimating head motion parameters; (6) registering functional images to high resolution T1 images using boundary-based registration; (7) removing nuisance factors such as head motion, CSF and white matter signals using ICA-AROMA (85); (8) removing linear and quadratic trends of the time series; (9) projecting volumetric time series to surface space (the *fsaverage5* model with medial wall masked out); (10) 6mm spatial smoothing. All preprocessing scripts are publicly available on github (https://github.com/zuoxinian/CCS). Any resting-state scan with a mean head motion above 0.5 mm was excluded from further analysis. The demographic information of subjects included in the analyses is listed in Supplemental Table 2.

### Gradient Analysis

Functional connectivity (FC) matrix and the corresponding Fisher-z transformed values were first generated for each resting-scan per visit. Then the two test-retest FC fisher-z matrices within one visit were averaged to increase signal-to-noise ratio for generating individual FCz matrix, which was later averaged across individuals to form group-level FCz matrix. For group-level FCz matrix, only the top 10% connections of each vertex were retained, other elements in the matrix were set to 0 to enforce sparsity. We then calculated the cosine distance between any two rows of the FCz matrix and subtracted from 1 to obtain a symmetrical similarity matrix.

Diffusion map embedding (27, 54) was implemented on the similarity matrix to derive gradients (https://github.com/NeuroanatomyAndConnectivity/gradient_analysis). The gradients were ordered by the variance explained (Supplemental Figure 1). To determine the corresponding relationship of the gradients across age groups we calculated the Pearson correlation coefficient between any two pairs of gradient maps across 1 year-interval age groups (Figure 2A). The gradient maps were first concatenated within gradients across age groups, for example gradient 1 maps in each age group were concatenated to form a matrix, then the gradient matrices were combined to generate the final global gradient matrix. The gradient maps are row vectors in the final matrix, and ordered by age, i.e. the first 11 rows are the first gradient maps from age 7 to age 17.

Gradient maps of Children and Adolescents group are summarized in Figure 1 and 2. The first 6 gradient maps across 1 year-interval age groups are listed in Figure 3. To quantify the gradient transitions across age groups, we extracted the gradient z values within default network regions as summarized through the Yeo et al., (3) 7-network solution averaged across the 400-parcel functional atlas of Schaefer and colleagues ((60); Figure 3B). To illustrate the gradient-level maturation of the default network, we averaged the maturation values within boundaries of seven functional networks in six gradient maturation maps, and summarized them in radar maps for each network (Figure 3C; Supplemental Figure 2). In each radar map, the six dimensions refer to the averaged maturation ages for each gradient. Box plots in Figure 4 display the manner in which gradients values are distributed across large-scale brain networks, ordered by the median value within each network.

### Maturation Map

Data from the 6- and 7-year-old participants was used to set the gradient orders. First, the six gradient maps were aligned according to their correlation coefficients to ensure gradient maps representing similar functional architectures across age groups. Here, the first gradient reflects a unimodal architecture, while the second gradient exhibits the differentiation of unimodal and transmodal regions. Then the maps were normalized by computing the standard z-scores across all surface vertices. For each vertex, the age at which the standard gradient value reaches its peak was assigned to this vertex to be its ‘maturation age’ (Figure 5A). In total, we generated six maturation maps for six gradient maps. The average of these maturation maps was obtained to represent the process of functional maturation. To explore whether this maturation map follows the order of cognitive processing hierarchy, we average the maturation values within boundaries of the 7 functional networks (3), as visualized in the Figure 5B bar graph. The corresponding error bars denote standard deviations across vertices.

## Acknowledgements

This work was supported by the Beijing Municipal Science and Tech Commission (Grant Z161100002616023 to X.N.Z), the Natural Science Foundation of China (Grant 81220108014 to X.N.Z), the China - Netherlands CAS-NWO Programme (Grant 153111KYSB20160020 to X.N.Z), the National R&D Infrastructure and Facility Development Program of China, Fundamental Science Data Sharing Platform (Grant DKA2020-12-02-21 to X.N.Z), the Startup Funds for Leading Talents at Beijing Normal University, Guangxi BaGui Scholarship (Grant 201621 to X.N.Z) and the National Institutes of Health (Grant R01MH120080 to A.J.H).

## Supplementary Material

**Supplemental Table 1.**
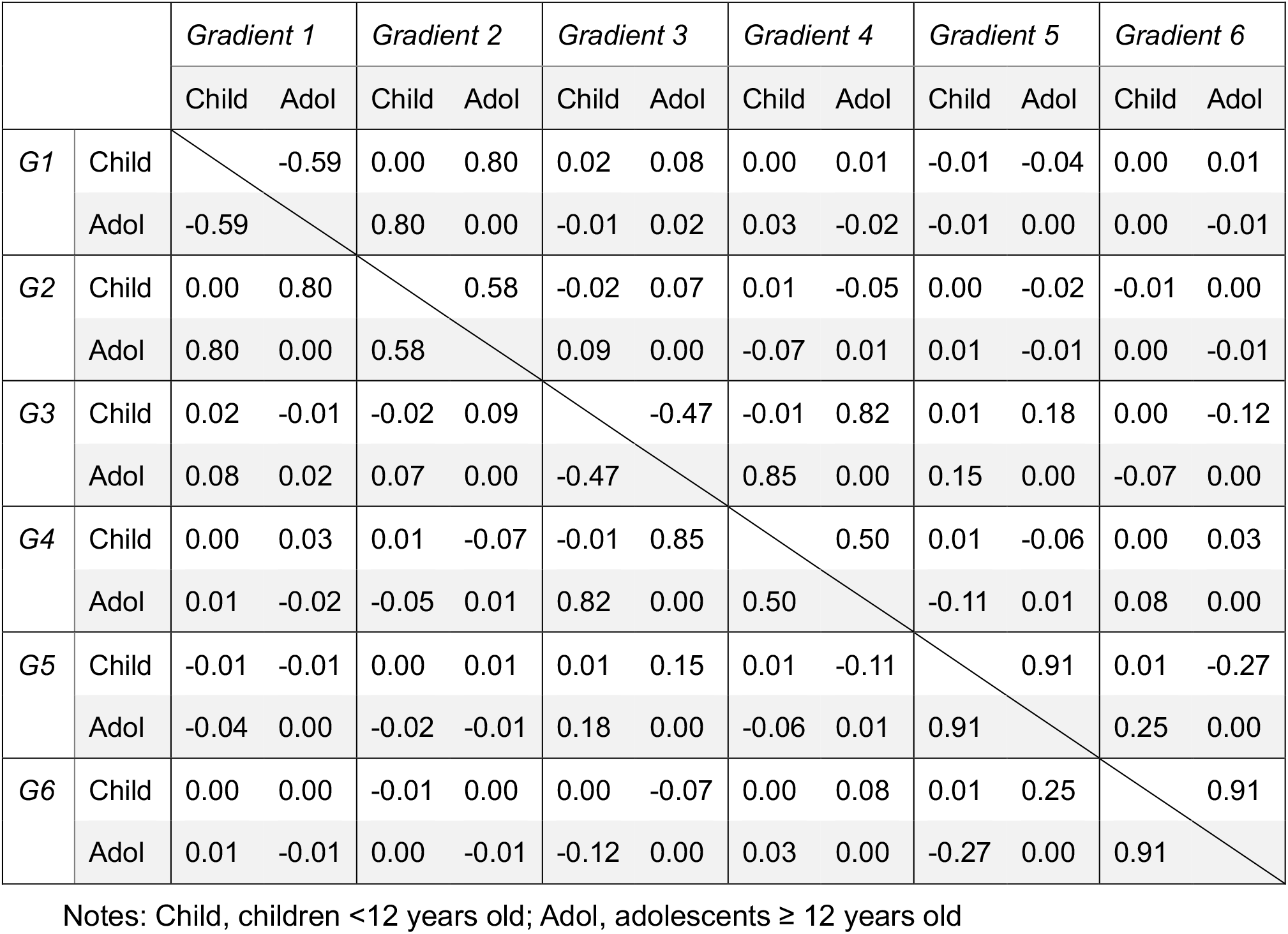
Pearson correlations for the first six gradients across children and adolescents.

|  |  | *Gradient 1* |  | *Gradient 2* |  | *Gradient 3* |  | *Gradient 4* |  | *Gradient 5* |  | *Gradient 6* |  |
| --- | --- | --- | --- | --- | --- | --- | --- | --- | --- | --- | --- | --- | --- |
|  |  | Child | Adol | Child | Adol | Child | Adol | Child | Adol | Child | Adol | Child | Adol |
| G1 | Child |  | -0.59 | 0.00 | 0.80 | 0.02 | 0.08 | 0.00 | 0.01 | -0.01 | -0.04 | 0.00 | 0.01 |
|  | Adol | -0.59 |  | 0.80 | 0.00 | -0.01 | 0.02 | 0.03 | -0.02 | -0.01 | 0.00 | 0.00 | -0.01 |
| G2 | Child | 0.00 | 0.80 |  | 0.58 | -0.02 | 0.07 | 0.01 | -0.05 | 0.00 | -0.02 | -0.01 | 0.00 |
|  | Adol | 0.80 | 0.00 | 0.58 |  | 0.09 | 0.00 | -0.07 | 0.01 | 0.01 | -0.01 | 0.00 | -0.01 |
| G3 | Child | 0.02 | -0.01 | -0.02 | 0.09 |  | -0.47 | -0.01 | 0.82 | 0.01 | 0.18 | 0.00 | -0.12 |
|  | Adol | 0.08 | 0.02 | 0.07 | 0.00 | -0.47 |  | 0.85 | 0.00 | 0.15 | 0.00 | -0.07 | 0.00 |
| G4 | Child | 0.00 | 0.03 | 0.01 | -0.07 | -0.01 | 0.85 |  | 0.50 | 0.01 | -0.06 | 0.00 | 0.03 |
|  | Adol | 0.01 | -0.02 | -0.05 | 0.01 | 0.82 | 0.00 | 0.50 |  | -0.11 | 0.01 | 0.08 | 0.00 |
| G5 | Child | -0.01 | -0.01 | 0.00 | 0.01 | 0.01 | 0.15 | 0.01 | -0.11 |  | 0.91 | 0.01 | -0.27 |
|  | Adol | -0.04 | 0.00 | -0.02 | -0.01 | 0.18 | 0.00 | -0.06 | 0.01 | 0.91 |  | 0.25 | 0.00 |
| G6 | Child | 0.00 | 0.00 | -0.01 | 0.00 | 0.00 | -0.07 | 0.00 | 0.08 | 0.01 | 0.25 |  | 0.91 |
|  | Adol | 0.01 | -0.01 | 0.00 | -0.01 | -0.12 | 0.00 | 0.03 | 0.00 | -0.27 | 0.00 | 0.91 |  |

Notes: Child, children <12 years old; Adol, adolescents ≥ 12 years old

**Supplemental Table 2.** Age and sex composition in CCNP sample

| Age Range | Percent Female | Total N |
| --- | --- | --- |
| 6.0-7.9 | 50 | 24 |
| 8.0-8.9 | 58 | 26 |
| 9.0-9.9 | 42 | 36 |
| 10.0-10.9 | 59 | 61 |
| 11.0-11.9 | 40 | 55 |
| 12.0-12.9 | 46 | 39 |
| 13.0-13.9 | 58 | 36 |
| 14.0-14.9 | 41 | 29 |
| 15.0-15.9 | 77 | 22 |
| 16.0-16.9 | 77 | 30 |
| 17.0-17.9 | 75 | 20 |
| Total Sample | 54 | 378 |
| Children (<12) | 40 | 202 |
| Adolescents (≥12) | 60 | 176 |

**Supplemental Figure 1:**
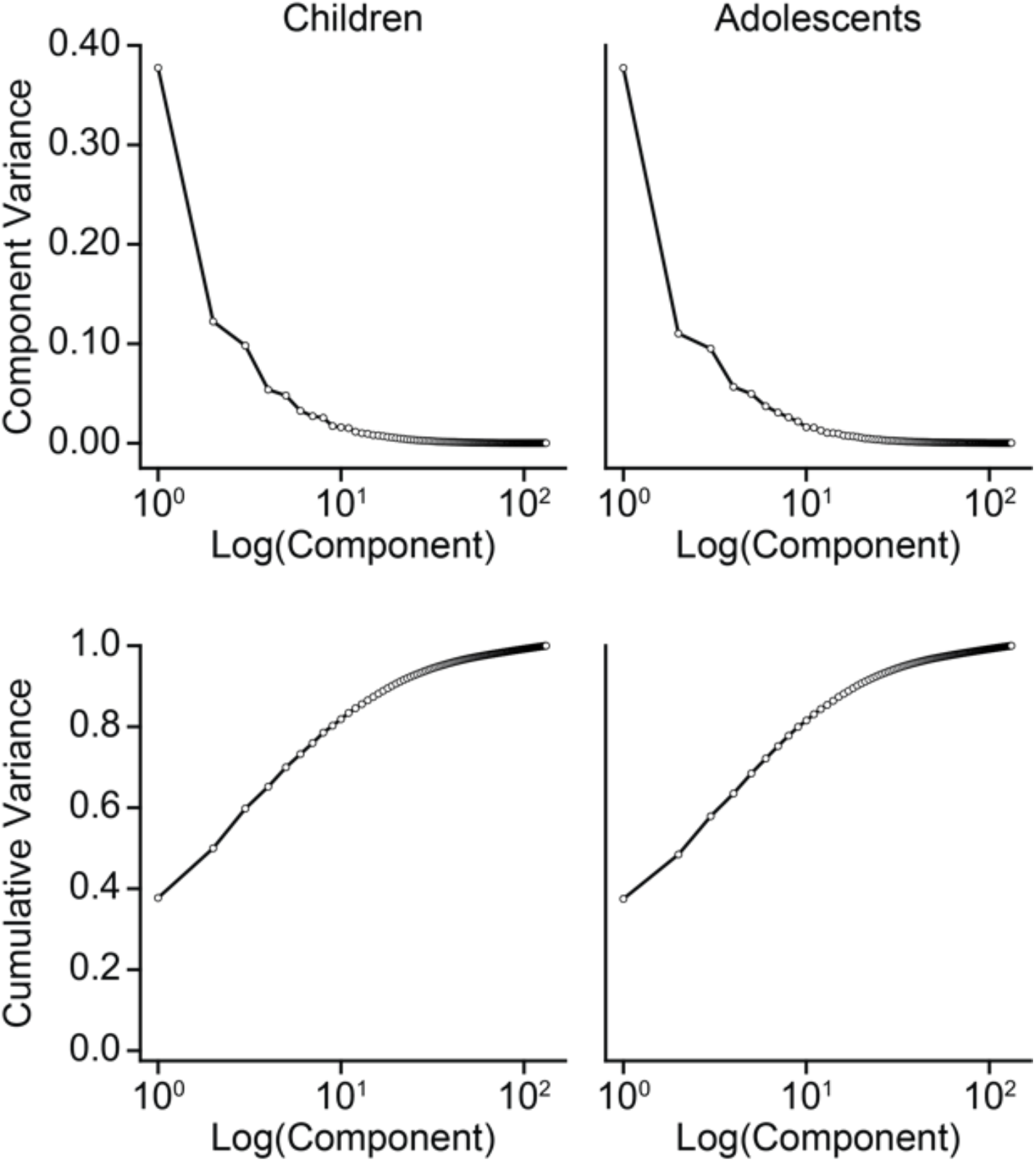
Variance (up row Y axis) and cumulative variance (bottom row Y axis) accounted by gradient components in Child (left column) and Adolescent (right column) groups. X axis reflects the logarithm of the gradient components number. The first gradient accounts for 37.7% and 37.5% variance in children and adolescents group respectively, the cumulative variance accounted by the first 6 gradients is 73.2% in children and 72.2% in adolescents.

**Supplemental Figure 2:**
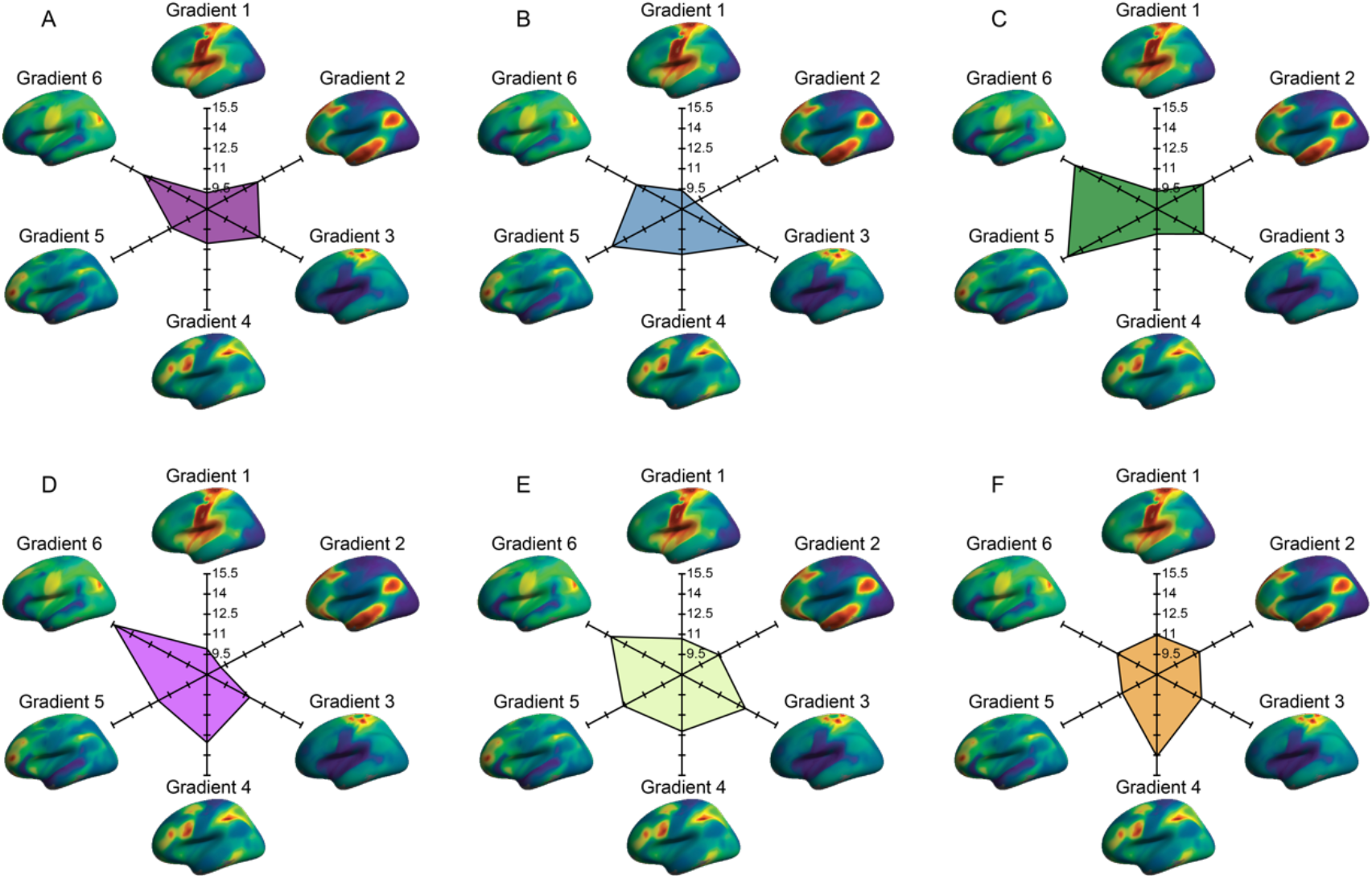
Radar plot displays the age at which network gradient values peak during development across each of the first six gradients. Network values for the (A) visual, (B) somato/motor, (C) salience/ventral attention, (D) dorsal attention, (E) limbic, and (F) frontoparietal control networks were summarized based on the Yeo et al., (2011) 7-network solution averaged across the 400-parcel functional atlas of Schaefer and colleagues (2018).

